# RNA-Seq analysis of compatible and incompatible styles of *Pyrus* species at the beginning of pollination

**DOI:** 10.1101/344044

**Authors:** Yaqin Guan, Kun Li, Yongzhang Wang, Chunhui Ma

## Abstract

In Rosaceae, incompatible pollen can penetrate into the style during the gametophytic self-incompatibility response. It is therefore considered a stylar event rather than a stigmatic event. In this study, we explored the differences in gene expression between compatibility and incompatibility in the early stage of pollination. The self-compatible pear variety “Jinzhuili” is a naturally occurring bud mutant from “Yali”, a leading Chinese native cultivar exhibiting typical gametophytic self-incompatibility. We collected the styles of ‘Yali’ and ‘Jinzhuili’ at 0.5 and 2 h after self-pollination and then performed high-throughput sequencing. According to the pathway enrichment analysis of the differentially expressed genes, “plant-pathogen interaction” was the most represented pathway. Quantitative PCR was used to validate these differential genes. The expression levels of genes related to pollen growth and disease inhibition, such as LRR (LEUCINE-RICH REPEAT EXTENSIN), resistance, and defensin, differed significantly between compatible and incompatible pollination. Interestingly, at 0.5 h, most of these genes were upregulated in the compatible pollination system compared with the incompatible pollination system. Calcium ion transport, which requires ATPase, also demonstrated upregulated expression. In summary, the self-incompatibility reaction was initiated when the pollen came into contact with the stigma.

## Introduction

Self-incompatibility (SI) is the general term used to describe several genetic mechanisms in angiosperms that prevent self-fertilization and thus encourage outcrossing and allogamy. RNase-based gametophytic self-incompatibility (GSI) is the most phylogenetically widespread, occurring in at least three families: Solanaceae, Rosaceae, and Scrophulariaceae (Igic *et al.*, 2004). Most fruit tree species belong to family Rosaceae and subfamily Spiraeoideae, which includes the supertribe Spiraeoideae that consists of the tribes Maleae (e.g., *Malus* and *Pyrus*) and Amygdaleae (e.g., *Prunus*), all of which exhibit typical gametophytic self-incompatibility (Shulaev *et al.*, 2008). Wu et al. (2012) reviewed the research progress in GSI mechanisms in fruit crops such as pear, apple, almond, sweet cherry, apricot, Japanese apricot, plum, and sour cherry (Wu *et al.*, 2012). As pear was the first fruit tree species in which S-ribonucleases were identified, the complete history of S-genotyping and technical developments can be monitored by evaluating the reported experiments (Halász & Hegedûs, 2006). The draft genome of pear (*Pyrus bretschneideri*) was generated using a combination of BAC-by-BAC and next-generation sequencing (Wu *et al.*, 2013b).

In Rosaceae, GSI is controlled by a single multi-allelic S locus that is composed of the pistil-S and pollen-S genes. The pistil-S gene encodes a polymorphic ribonuclease (S-RNase) essential for discriminating self-pollen; however, the S-RNase system has not been fully elucidated. SLF (S-locus F-box) genes appear to be involved in the determination of pollen-S specificity, but the function of F-box genes in the SI reaction remains unknown. There is no direct evidence of the interaction between SLF and different alleles/domains of the S-RNase protein. McClure (2006) proposed the compartmentalization model of GSI, but no evidence exists regarding when the vesicles begin to break down following pollination incompatibility (McClure, 2006). In addition, a number of researchers suggested that S-RNase specifically induces tip-localized reactive oxygen species (ROS) disruption, actin cytoskeleton depolymerisation, and nuclear DNA degradation in the incompatible pollen tubes of pear (Liu *et al.*, 2007; Wang *et al.*, 2010a). We also found that self S-RNase selectively blocks the activity of Ca^2+^ channels at the apical pollen tube of pear (Qu *et al.*, 2016). It has been suggested that pollen rejection and incompatibility systems are interconnected and should be regarded as different aspects of a single system that provide fine control over plant fertilization (Cruz-Garcia *et al.*, 2003).

For fruit trees, including pear, a stable transgenic system is highly difficult to construct. However, we were able to obtain ‘Jinzhuili’, which is a naturally occurring self-compatible mutant of ‘Yali’ (*Pyrus bretschneideri* Rehd.). Field pollination tests showed that >76% of flowers set fruit and many pollen tubes grew down to the base of the style during ‘Jinzhuili’ self-pollination and cross-pollination of ‘Yali’ (female) × ‘Jinzhuili’ (male). In contrast, fruit setting did not occur and most pollen tubes were arrested at the upper part of the style during ‘Yali’ self-pollination and cross-pollination of ‘Jinzhuili’ (female) × ‘Yali’ (male). These results indicate that ‘Yali’ and ‘Jinzhuili’ exhibit a normal SI response in the style, rejecting competent ‘Yali’ pollen but accepting ‘Jinzhuili’ pollen. Further molecular analyses revealed that both ‘Yali’- and ‘Jinzhuili’-labelled S21-RNase and S34-RNase genes were transcribed normally with identical mRNA sequences. Therefore, we suggest that ‘Jinzhuili’ has most likely lost pollen SI activity, resulting in a style that behaves normally during the SI response, similar to that of the wild-type ‘Yali’. However, Wu et al. (2013) identified five SLF (S-Locus F-box) genes in ‘Yali’ but no nucleotide differences were observed in the SLF genes of ‘Jinzhuili’ (Wu *et al.*, 2013a). We selected ‘Jinzhuili’ and ‘Yali’, two cultivars of the same species with the same genetic background but with some differing genetic mutations, to perform high-throughput sequencing in order to explore the variation in gene expression between compatible and incompatible pollination.

Present research on self-incompatibility in Rosaceae has predominantly focused on the response of the pollen tubes in the style, but little data exists regarding the role of the stigma at the start of pollination. Therefore, we selected the styles as the tissue with which to perform high-throughput sequencing analysis at 0.5 h and 2 h after pollination.

## Results

### *De novo* assembled transcriptomes of self-compatible and self-incompatible styles

Five whole-plant RNA bulks derived from the styles of ‘Jinzhuili’ at 0.5 h (JZ_0.5h) and 2 h (JZ_2h) after self-pollination and the styles of ‘Yali’ without pollination (control, YL_CK), and at 0.5 h (YL_0.5h) and 2 h (YL_2h) after self-pollination were utilized for transcriptome sequencing analysis to identify genes that were differentially expressed between the self-compatible and self-incompatible style and the pollen tube. RNA sequencing (RNA-Seq) generated approximately 4.46 Gb raw reads from each of the five transcriptomes. After filtering the reads, the clean reads were mapped to the reference genome using HISAT (Kim *et al.*, 2015). An average of 59.32% reads was mapped, and the uniformity of the mapping result for each sample suggested that the samples were comparable. The mapping details are shown in Table 1. After mapping the sequenced reads to the reference genome and reconstructing the transcripts, we ultimately obtained 8,559 novel transcripts from all of the samples, of which 5,768 constitute previously unknown splicing events for a known gene; 1,185 are novel coding transcripts without any known features; and the remaining 1,606 are long noncoding RNAs.

**Table 1.** Assembled and annotated transcriptomes from self-compatible and self-incompatible styles (JZ_0.5, YL_0.5, JZ_2 and YL_2, respectively) and styles without pollination as control.

| Sample | Total Raw Reads (Mb) | Total Clean Reads (Mb) | Total Clean Bases (Gb) | Clean Reads Q20(%) | Clean Reads Q30 (%) | Clean Reads Ratio (%) |
| --- | --- | --- | --- | --- | --- | --- |
| JZ_0.5 | 35.93 | 30.14 | 4.52 | 98.47 | 95.25 | 83.87 |
| JZ_2 | 35.05 | 29.91 | 4.49 | 98.85 | 96.36 | 85.31 |
| JZ_CK | 35.05 | 29.73 | 4.46 | 98.87 | 96.42 | 84.82 |
| YL_0.5 | 35.93 | 29.41 | 4.41 | 98.50 | 95.31 | 81.87 |
| YL_2 | 35.05 | 29.55 | 4.43 | 98.96 | 96.61 | 84.29 |

Note: (1) Q20 indicate the rate of bases which quality is greater than 20. (2) YL\_0.5 and YL\_2 is that the styles of ‘Yali’ were collected at 0.5 and 2 h after self-pollination. JZ\_0.5 and JZ\_2 is that the styles of ‘Jinzhuli’ were collected at 2 h after self-pollination. JZ\_CK is that styles of ‘Jinzhuli’ without pollination as control.

### Differential expression statistics

There were 1,795 and 1,658 differentially expressed genes (DEGs) between the style of the control and ‘Jinzhuili’ at 0.5 and 2 h after self-pollination, respectively. There were 3,842 and 3,787 DEGs between the style of the control and the style of ‘Yali’ at 0.5 and 2 h following self-pollination, respectively. At 0.5 and 2 h after self-pollination, there were 3,453 and 3,212 DEGs between the style of ‘Jinzhuili’ and ‘Yali’, respectively (Figure 1). Within 2 h after self-pollination, the differences in gene expression between the compatible style and incompatible style had begun to decrease. Following compatible or incompatible pollination, at which point the pollen tube was growing in the style, the differences in gene expression between the style and the control style decreased progressively. The number of genes expressed in the style after incompatible pollination was higher than that in the style after compatible pollination. This result suggests that gene expression is more critical at the moment that the pollen comes into contact with the stigma. Furthermore, the results suggest that gene expression is more complex under incompatible pollination.

**Figure 1.**
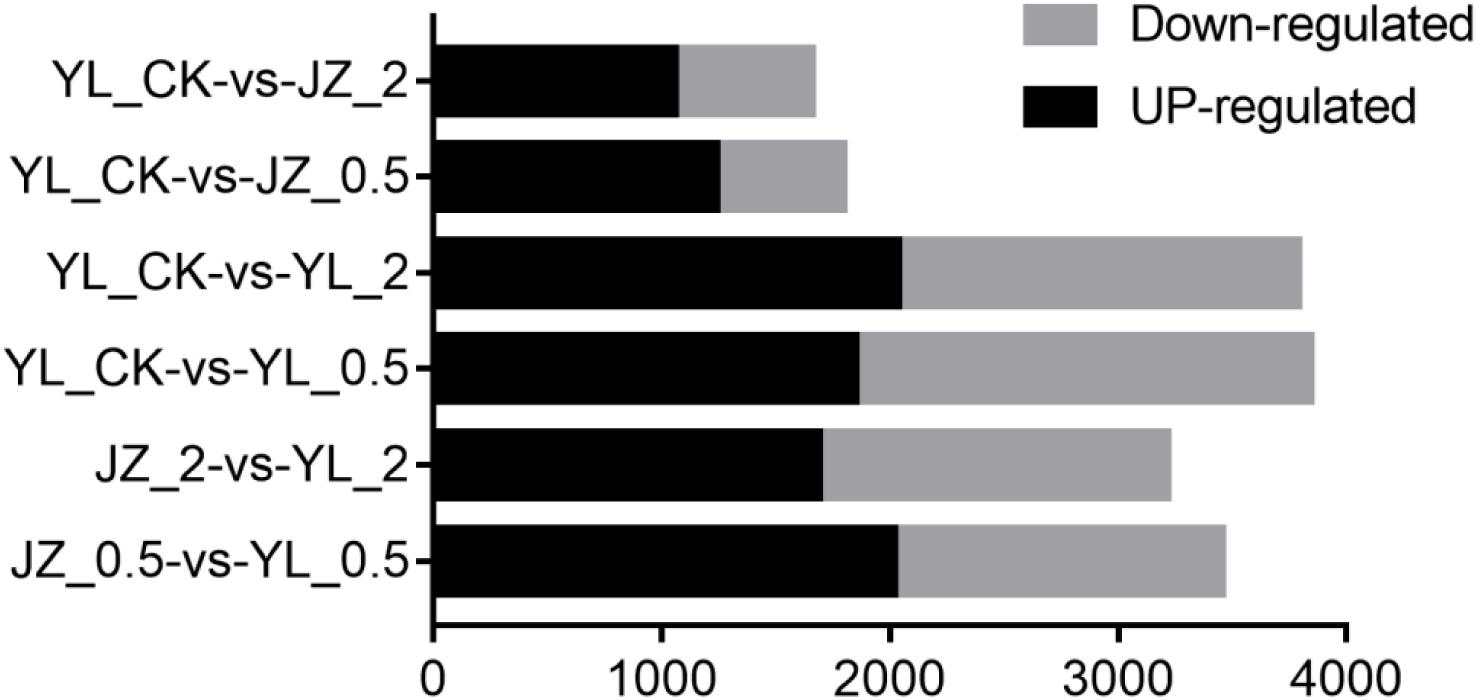
Summary of DEGs (Differentially Expressed Gene). Y axis represents comparing samples. X axis represents DEG numbers. Fold Change >= 2.00 and FDR <= 0.001.

#### Kyoto Encyclopedia of Genes and Genomes (KEGG) enrichment analysis of DEGs

KEGG (http://www.genome.jp/kegg/) is a knowledge base for the systematic analysis of gene functions in terms of the networks of genes and molecules. The process of the pollen falling away from the stamens and then falling onto the stigma during fertilization is a highly complex process that includes a series of events. Enrichment analysis was used to initially characterize the style transcriptome after pollination with regards to various biological activities that were sorted by relevance, as well as to identify genes that were differentially expressed between the compatible and incompatible plants. At 0.5 h and 2 h, the DEG terms involved in “plant-pathogen interaction”, “Protein processing in endoplasmic reticulum”, “Pentose and glucuronate interconversions” after compatible pollination (Figure 2A and B), but the DEG terms are mainly came from “Protein processing in endoplasmic reticulum”, “glycolysis/gluconeogenesis”, “glycosphingolipid biosynthesis-ganglio” series after incompatible pollination (Figure 2A and C). Furthermore, we calculated the Pearson’s correlation coefficient between all samples based on gene expression, as indicated in Figure 3. At 0.5 h, compatible pollination (JZ_0.5h) was not correlated with incompatible pollination (YL_0.5h) in terms of gene expression, whereas by 2 h the correlation coefficient had increased. These results suggest that the mutual recognition of pollen and the stigma and the incompatibility (compatibility) reaction was initiated when the pollen had fallen onto the stigma.

**Figure 2.**
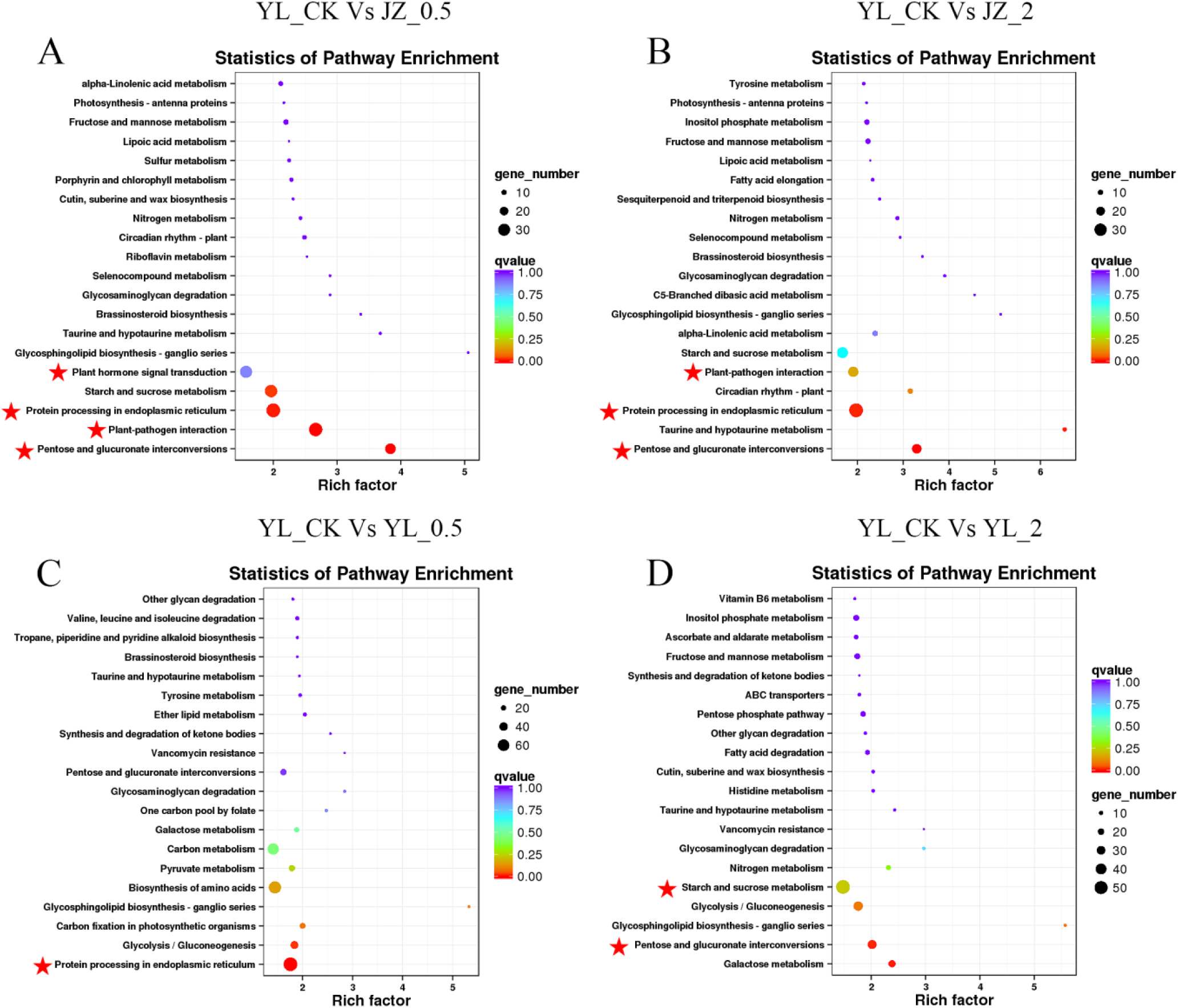
Pathway functional enrichment of DEGs. (A) Enriched pathways in self-compatible at 0.5 h (YL_CK Vs JZ_0.5). (B) Enriched pathways in self-compatible at 2 h (YL_CK Vs JZ_2). (C) Enriched pathways in self-incompatible at 0.5 h (YL_CK Vs YL_0.5). (D) Enriched pathways in self-incompatible at 2 h (YL_CK Vs YL_2). X axis represents enrichment factor. Y axis represents pathway name. Coloring indicate qvalue (high: violet, low: red), the lower qvalue indicate the more significant enriched. Pointsize indicate DEG number (more: big, less: small). Pentagram shows the significantly enriched pathway.

**Figure 3.**
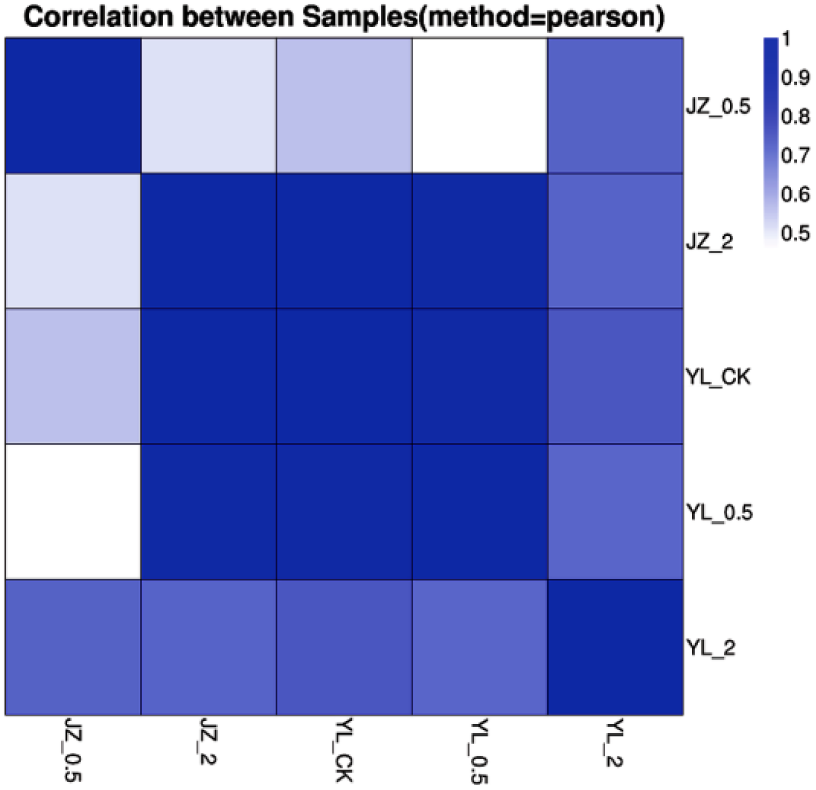
Heatmap of pearson correlation between samples. Both X and Y axis represent each sample. Coloring indicate pearson correlation (high: blue, low: white). YL_0.5 and YL_2 is that the styles of ‘Yali’ were collected at 0.5 and 2 h after self-pollination. YL_CK is that ‘Yali’ without pollination as control. JZ_0.5 and JZ_2 is that the styles of ‘Jinzhuili’ were collected at 2 h after self-pollination.

#### Pollen interaction with the stigma

The communication of changes in the extracellular matrix to the interior of the cell is crucial for cell functioning (Gao *et al.*, 2014). *Arabidopsis* RALF4 and RALF19 (Rapid alkalinization factors) regulate pollen tube integrity and growth, and their function depends on the pollen-expressed proteins of the LEUCINE-RICH REPEAT EXTENSIN (LRX) family (Mecchia *et al.*, 2017). LRR (Leu-rich repeat) proteins are involved in specific protein–protein interactions and are confined predominantly to eukaryotes. The LRR proteins have a significant role in plant defense. Resistance to a diverse range of pathogens, including nematodes, fungi, bacteria, and viruses, involves LRR proteins either as resistance proteins or as proteins required for the functioning of resistance proteins. These proteins might be involved in the defense of reproductive tissues from pathogens or may be involved in reproduction itself (Jones & Jdg, 1997; Ndinyanka *et al.*, 2017). Pollen–stigma interactions are similar to host-pathogen interactions (Kovaleva & Zakharova, 2003; Mondragon *et al.*, 2017; Wilkins *et al.*, 2014). Pollen self-incompatibility and species incompatibility may also be viewed as forms of defense. It would not be surprising if the parallels between resistance and pollen incompatibility extended to the involvement of LRR proteins in both processes. We searched for pollen-related DEGs (*P*≤0.05) in JZ_0.5-VS-YL_0.5 and JZ_2-VS-YL_2. We discovered 13 pollen-specific DEGs, 12 of which were related to the LRR gene (Table 2). We selected these 13 genes for quantitative expression analysis in the style following pollination at different times. The quantitative PCR and RNA-Seq results were identical at 0.5 h, whereas at 2 h, only 38.46% of the results were consistent (Figure 4). At 0.5 h, all pollen-specific gene expression was significantly higher in compatible pollination than in incompatible pollination. However, at 2 h, 13 pollen-specific genes were significantly more highly expressed in incompatible pollination than in compatible pollination, of which nine were significantly different (*P*<0.01). We counted the DEGs of all LRXs at different time periods after pollination. The upregulated expression ratio of this gene was 83.61% at 0.5 h, but had declined to 41.38% at 2 h (Figure 5). These results suggested that following pollen and stigma identification, the regulation of pollen tube growth under the condition of compatible pollination occurs earlier than that under incompatible pollination.

**Table 2.** The differences in the expression of 13 pollen-specific genes and their definitions.

| GeneID | $\text{Log}_2^{(\text{JZ\_0.5/YL\_0.5})}$ | $\text{Log}_2^{(\text{JZ\_2/YL\_2})}$ | Definition |
| --- | --- | --- | --- |
| LOC103927207 | 2.16 | -3.33 | PREDICTED: pollen-specific leucine-rich repeat extensin-like protein 1 [*Pyrus x bretschneideri*] |
| LOC103932328 | 1.55 | -3.68 | PREDICTED: pollen-specific protein SF3-like [*Malus domestica*] |
| LOC103943315 | 1.93 | -1.03 | PREDICTED: pollen-specific leucine-rich repeat extensin-like protein 3 [*Pyrus x bretschneideri*] |
| BGI_novel_G001157 | 6.80 | -1.30 | PREDICTED: pollen-specific leucine-rich repeat extensin-like protein 4 [*Pyrus x bretschneideri*] |
| BGI_novel_G001158 | 3.33 | -1.30 | PREDICTED: pollen-specific leucine-rich repeat extensin-like protein 4 [*Pyrus x bretschneideri*] |
| LOC103934404 | 3.60 | 3.89 | PREDICTED: pollen-specific leucine-rich repeat extensin-like protein 4 [*Pyrus x bretschneideri*] |
| LOC103937306 | 1.90 | -2.53 | PREDICTED: pollen-specific leucine-rich repeat extensin-like protein 3 [*Pyrus x bretschneideri*] |
| LOC103942292 | 1.23 | 1.21 | PREDICTED: pollen-specific leucine-rich repeat extensin-like protein 3 [*Pyrus x bretschneideri*] |
| LOC103942293 | 1.23 | 1.21 | PREDICTED: pollen-specific leucine-rich repeat extensin-like protein 3 [*Pyrus x bretschneideri*] |
| LOC103942771 | 1.81 | 1.77 | PREDICTED: pollen-specific leucine-rich repeat extensin-like protein 4 [*Pyrus x bretschneideri*] |
| BGI_novel_G001159 | 4.33 | 3.17 | PREDICTED: pollen-specific leucine-rich repeat extensin-like protein 4 [*Pyrus x bretschneideri*] |
| BGI_novel_G001162 | 2.16 | 2.08 | PREDICTED: pollen-specific leucine-rich repeat extensin-like protein 4 [*Pyrus x bretschneideri*] |
| BGI_novel_G001163 | 2.17 | 1.81 | PREDICTED: pollen-specific leucine-rich repeat extensin-like protein 4 [*Pyrus x bretschneideri*] |

Note:
(1) YL\_0.5 and YL\_2 is that the styles of ‘Yali’ were collected at 0.5 and 2 h after self-pollination. JZ\_0.5 and JZ\_2 is that the styles of ‘Jinzhuli’ were collected at 2 h after self-pollination.
(2) Positive values represent up-regulated gene and Negative values represent the down regulated genes. FDR-corrected $P \leq 0.0005$

**Figure 4.**
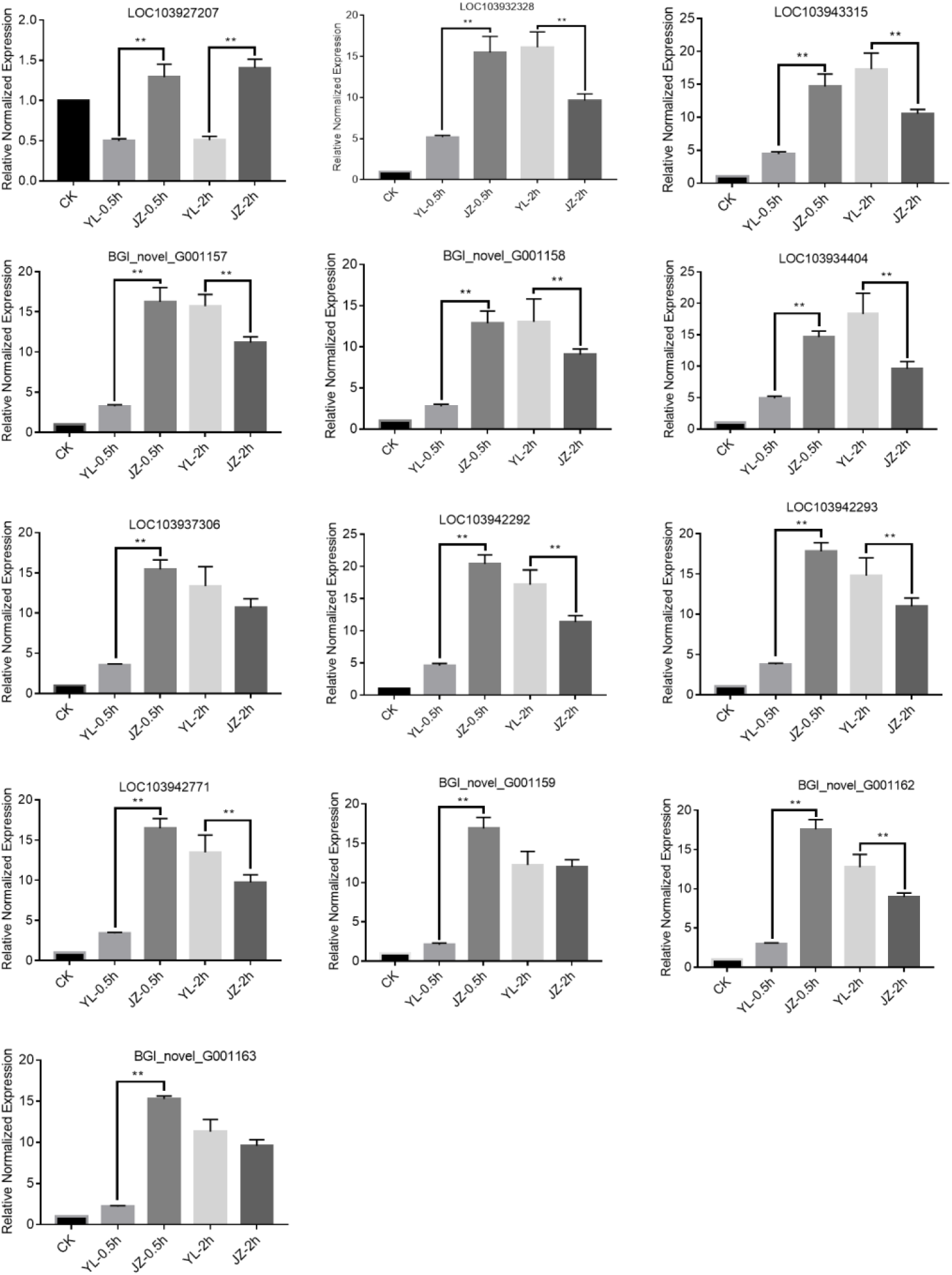
Expression of 13 pollen-specific genes in styles measured by quantitative PCR. Comparison of the expression of 13 pollen-specific genes during compatible and incompatible pollination. The 13 genes were selected according to the high-throughput sequencing results. ‘Yali’ styles were collected 0.5 and 2 h after self-pollination (YL_0.5h and YL_2h). ‘Jinzhuili’ styles were collected 0.5 and 2 h after self-pollination (JZ_0.5h and JZ_2h). Non-pollinated ‘Yali’ styles were collected after 0.5 h without pollination as the control (YL_CK). The data are presented as the means ± standard error. ** Highly significant data (P < 0.01).

**Figure 5.**
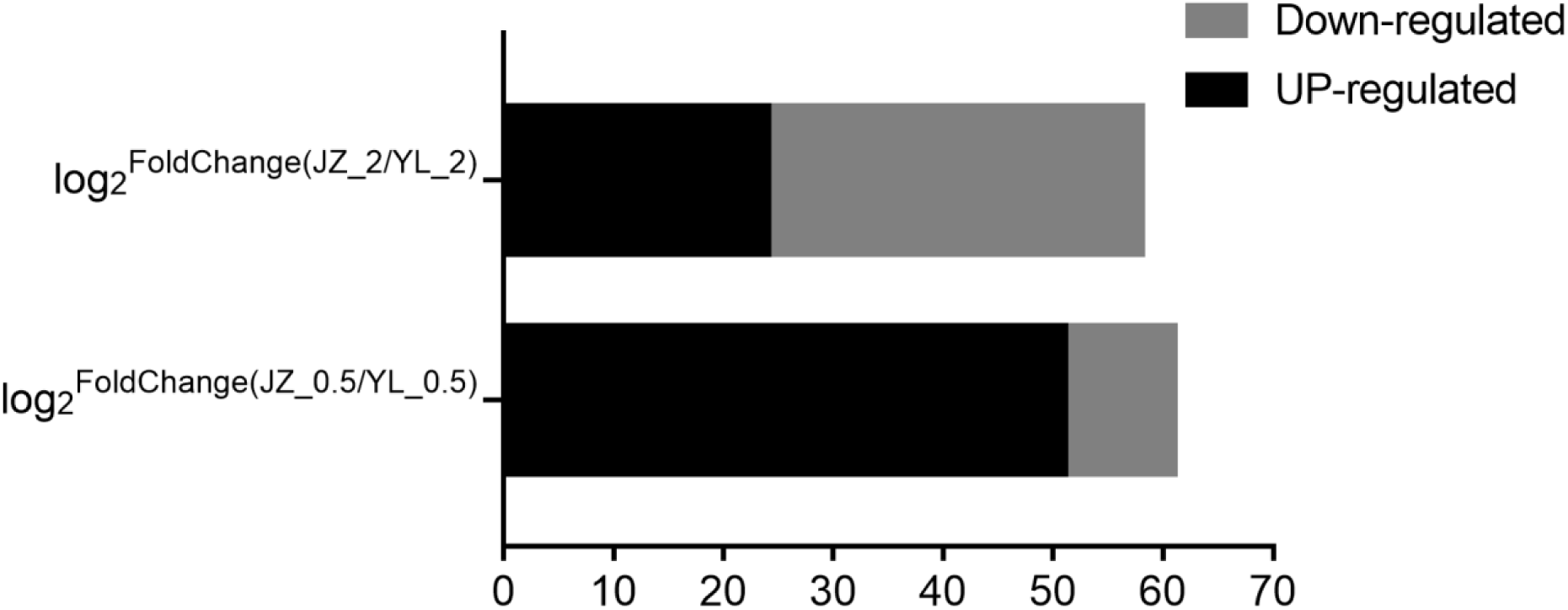
Summary of DEGs about leucine-rich repeat extensin-like protein. Y axis represents comparing samples. X axis represents DEG numbers. Fold Change >= 2.00 and FDR <= 0.001.

There were 17 disease resistance-related genes involved in the regulation of the defense response to fungus according to the high-throughput sequencing results. At 0.5 h, there were 13 genes in 17 disease resistance genes, and their quantitative PCR results were consistent with the results of the high-throughput sequencing, whereas at 2 h, the results of the 15 genes were identical (Table 3). There were 15 and 14 genes at 0.5 h and 2 h, respectively, that exhibited higher expression levels in the compatible pollination than in the incompatible pollination (Figure 6). The function of defensin-like proteins is in the defense response to fungus (http://www.uniprot.org/uniprot/P20159). There were eight genes associated with defensin-like proteins according to the high-throughput sequencing (Table 4). There were seven and six genes at 0.5 h and 2 h, respectively, that exhibited higher expression in compatible pollination than in incompatible pollination (Figure 7).

**Table 3.** The differences in the expression of 17 disease-resistance genes and their definitions.

| GeneID | $\text{Log}_2^{(\text{JZ}_{0.5}/\text{YL}_{0.5})}$ | $\text{Log}_2^{(\text{JZ}_{2}/\text{YL}_{2})}$ | Definition |
| --- | --- | --- | --- |
| LOC103930781 | -1.28 | 1.37 | PREDICTED: protein ENHANCED DISEASE RESISTANCE 2-like isoform X1 [*Pyrus x bretschneideri*] |
| LOC103936822 | 1.09 | -1.47 | PREDICTED: disease resistance RPP13-like protein 4 [*Pyrus x bretschneideri*] |
| LOC103937131 | 2.00 | 1.63 | PREDICTED: LOW QUALITY PROTEIN: disease resistance protein RPM1-like [*Malus domestica*] |
| LOC108865347 | 3.29 | -1.51 | PREDICTED: probable disease resistance protein At4g27220 isoform X1 [*Pyrus x bretschneideri*] |
| LOC103954680 | 7.82 | 9.02 | PREDICTED: probable disease resistance protein At4g27220 [*Malus domestica*] |
| LOC103956518 | 1.87 | 1.92 | PREDICTED: probable disease resistance protein At5g66900 [*Pyrus x bretschneideri*] |
| LOC103961932 | 10.76 | 10.75 | PREDICTED: disease resistance protein RPP8-like [*Pyrus x bretschneideri*] |
| LOC103961933 | 9.32 | 9.33 | PREDICTED: disease resistance protein RPP8-like [*Pyrus x bretschneideri*] |
| LOC103962513 | -1.21 | 1.95 | PREDICTED: disease resistance protein RPM1-like isoform X1 [*Pyrus x bretschneideri*] |
| LOC103941986 | 3.22 | 3.35 | PREDICTED: putative disease resistance protein RGA3 [*Pyrus x bretschneideri*] |
| LOC103956818 | -3.51 | -5.34 | PREDICTED: probable disease resistance protein At4g27220 [*Pyrus x bretschneideri*] |
| BGI_novel_G001126 | 3.96 | 4.06 | PREDICTED: putative disease resistance protein RGA4 [*Pyrus x bretschneideri*] |
| LOC103928191 | 1.65 | 1.08 | TIR-NBS-LRR disease resistance protein [*Malus domestica*] |
| LOC103949124 | -1.73 | 2.05 | PREDICTED: protein ENHANCED DISEASE RESISTANCE 2-like isoform X3 [*Pyrus x bretschneideri*] |
| LOC103949635 | 1.24 | 2.31 | PREDICTED: putative disease resistance protein At5g47280 [*Pyrus x bretschneideri*] |
| LOC103957587 | 5.31 | 3.35 | PREDICTED: probable disease resistance protein At5g66900 [*Malus domestica*] |
| LOC103961661 | -2.56 | -1.92 | PREDICTED: disease resistance protein RGA2-like [*Pyrus x bretschneideri*] |

Note:
(1) YL\_0.5 and YL\_2 is that the styles of ‘Yali’ were collected at 0.5 and 2 h after self-pollination. JZ\_0.5 and JZ\_2 is that the styles of ‘Jinzhuli’ were collected at 2 h after self-pollination.
(2) Positive values represent up-regulated gene and Negative values represent the down regulated genes. FDR-corrected $P \leq 0.0005$

**Figure 6.**
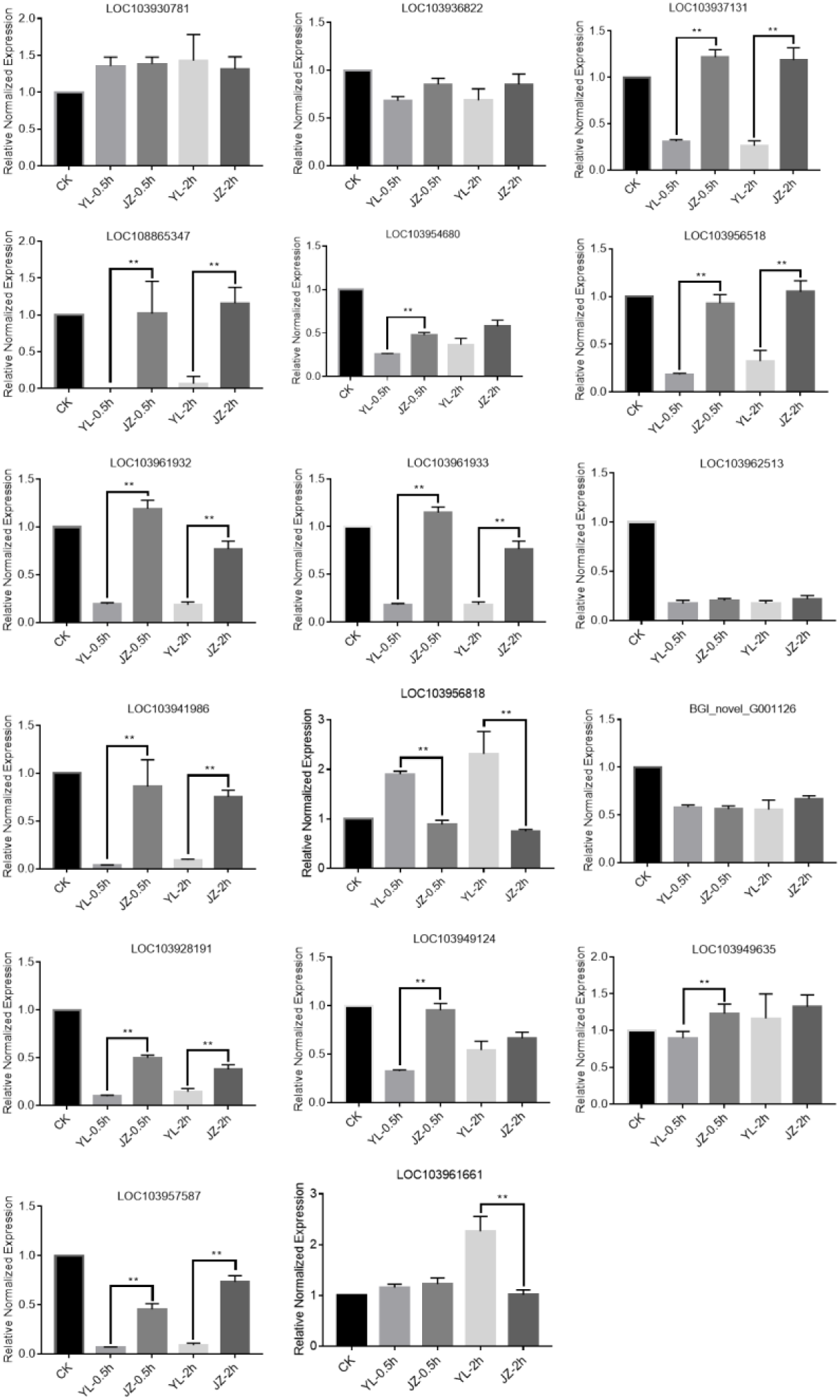
Expression of 17 disease-resistance genes in styles measured by quantitative PCR. Comparison of the expression of 17 disease-resistance genes during compatible and incompatible pollination. The 17 genes were selected according to the high-throughput sequencing results. ‘Yali’ styles were collected 0.5 and 2 h after self-pollination (YL_0.5h and YL_2h). ‘Jinzhuili’ styles were collected 0.5 and 2 h after self-pollination (JZ_0.5h and JZ_2h). Non-pollinated ‘Yali’ styles were collected after 0.5 h without pollination as the control (YL_CK). The data are presented as the means ± standard error. ** Highly significant data (P < 0.01).

**Table 4.** The differences in the expression of 8 defensin genes and their definitions.

| GeneID | $\text{Log}_2^{(\text{JZ}_{0.5}/\text{YL}_{0.5})}$ | $\text{Log}_2^{(\text{JZ}_2/\text{YL}_2)}$ | Definition |
| --- | --- | --- | --- |
| LOC103930708 | 1.15 | 1.53 | PREDICTED: defensin-like protein 2 [*Pyrus x bretschneideri*] |
| LOC103937764 | 2.01 | 2.64 | PREDICTED: defensin-like protein 2 [*Pyrus x bretschneideri*] |
| LOC103945528 | -1.24 | -1.96 | PREDICTED: defensin Ec-AMP-D2-like [*Malus domestica*] |
| LOC103943156 | 1.62 | -4.01 | PREDICTED: defensin Ec-AMP-D2-like [*Malus domestica*] |
| LOC103946497 | 1.13 | 1.35 | PREDICTED: defensin Tk-AMP-D1.1-like [*Pyrus x bretschneideri*] |
| LOC103946498 | 1.08 | 1.35 | PREDICTED: defensin Tk-AMP-D1.1-like [*Pyrus x bretschneideri*] |
| BGI_novel_G000142 | 2.04 | 2.84 | PREDICTED: defensin-like protein 2 [*Pyrus x bretschneideri*] |
| BGI_novel_G000791 | 2.02 | 1.81 | PREDICTED: defensin-like protein 2 [*Pyrus x bretschneideri*] |

Note:
(1) YL\_0.5 and YL\_2 is that the styles of ‘Yali’ were collected at 0.5 and 2 h after self-pollination. JZ\_0.5 and JZ\_2 is that the styles of ‘Jinzhuli’ were collected at 2 h after self-pollination.
(2) Positive values represent up-regulated gene and Negative values represent the down regulated genes. FDR-corrected $P \leq 0.0005$

**Figure 7.**
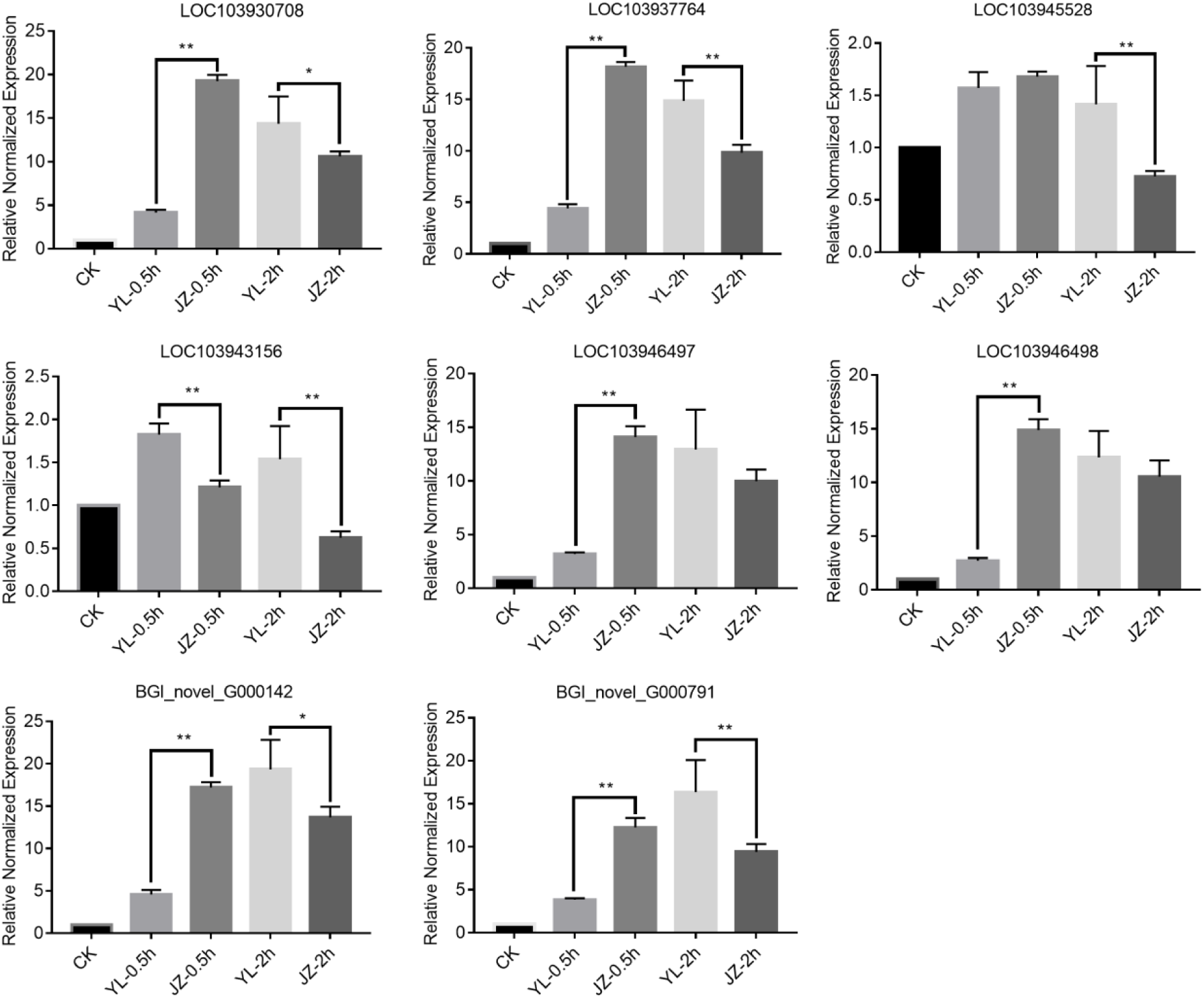
Expression of 8 defensin genes in styles measured by quantitative PCR. Comparison of the expression of 8 defensin genes during compatible and incompatible pollination. The 8 genes were selected according to the high-throughput sequencing results. ‘Yali’ styles were collected 0.5 and 2 h after self-pollination (YL_0.5h and YL_2h). ‘Jinzhuili’ styles were collected 0.5 and 2 h after self-pollination (JZ_0.5h and JZ_2h). Non-pollinated ‘Yali’ styles were collected after 0.5 h without pollination as the control (YL_CK). The data are presented as the means ± standard error. ** Highly significant data (P < 0.01).

The expression of these genes follows a trend; that is, at 0.5 h after pollination, the expression of most genes in the compatible pollination is higher than in the incompatible pollination. However, the expression of most of the pollen-specific and defensin-like protein genes differed at 2 h following pollination. Interestingly, there was a relatively high expression level of genes related to disease resistance at the beginning of the compatible pollination process. Hiscock and Mcinnis (2004) considered the sporophytic self-compatible response process of *Brassica* spp. to be similar to the host-pathogen (HP) interaction process. The transcription rate of genes for pathogenesis-related proteins was increased in only 5 min during the HP response (Hiscock & Mcinnis, 2004). In a previous study, after 48 h of pollination, the gene expression of pear (*P. bretschneideri*) was compared in the style, and it was speculated that the plants could identify compatible and incompatible pollen using the plant-pathogen signaling system at the start of pollination (Shi *et al.*, 2017). Therefore, we believe that the mutual recognition of the pollen and stigma in GSI of Rosaceae is similar to the pollen and stigma recognition process in the sporophytic self-incompatibility of *Brassica* spp.

#### ATPase

Little research on the role of ATPase in plant self-incompatibility exists. At present, the limited literature on the role of ATPases in self-incompatibility suggests that Ca^2+^-ATPase regulates the Ca^2+^ outflow of papilla cells or Ca^2+^ influx of the pollen tube on the stigma (Miao *et al.*, 2013). The tip of a pollen tube exhibiting normal growth is filled with a large number of mitochondria (Hill *et al.*, 2012), which may be related to the need for ATP during pollen tube growth. We selected nine differentially expressed ATPase-related genes (Table 5) according to the high-throughput sequencing results. After quantitative PCR verification, the expression of eight of the genes in the compatible pollination was higher than under incompatible pollination at 0.5 h following pollination (Figure 8), but only three of these genes were significantly different (*P*<0.01). These three genes are all associated with calcium transport (Figure 8). At 2 h after pollination, no regularity in the expression of ATPase related genes was observed. The results suggest that ATPase is involved in self-incompatibility through the calcium signalling pathway at the beginning of pollination.

**Table 5.** The differences in the expression of 10 genes about ATPase and their definitions

| GeneID | $\text{Log}_2^{(\text{JZ}_{0.5}/\text{YL}_{0.5})}$ | $\text{Log}_2^{(\text{JZ}_2/\text{YL}_2)}$ | Definition |
| --- | --- | --- | --- |
| LOC103926817 | 2.39 | 2.40 | PREDICTED: calcium-transporting ATPase 9, plasma membrane-type-like isoform X1 |
| LOC103927126 | 1.78 | -1.29 | PREDICTED: putative calcium-transporting ATPase 11, plasma membrane-type [*Pyrus x bretschneideri*] |
| LOC103929302 | 2.38 | 1.75 | PREDICTED: putative phospholipid-transporting ATPase 4 [*Pyrus x bretschneideri*] |
| LOC103932096 | -6.52 | -2.10 | PREDICTED: probable copper-transporting ATPase HMA5 [*Pyrus x bretschneideri*] |
| LOC103941070 | -3.20 | -2.11 | PREDICTED: 26S proteasome non-ATPase regulatory subunit 14 homolog [*Malus domestica*] |
| LOC103952641 | 2.19 | 1.83 | PREDICTED: calcium-transporting ATPase 2, plasma membrane-type-like [*Pyrus x bretschneideri*] |
| LOC103955945 | 3.38 | 4.19 | PREDICTED: ATPase 8, plasma membrane-type-like [*Pyrus x bretschneideri*] |
| LOC103961555 | 10.18 | 9.94 | PREDICTED: putative phospholipid-transporting ATPase 4 isoform X1 [*Pyrus x bretschneideri*] |
| LOC103962083 | -1.46 | -1.21 | PREDICTED: vacuolar proton ATPase a3-like [*Pyrus x bretschneideri*] |
| BGI_novel_G001178 | 4.20 | 2.98 | PREDICTED: ATPase WRNIP1-like [*Malus domestica*] |

Note:
(1) YL\_0.5 and YL\_2 is that the styles of ‘Yali’ were collected at 0.5 and 2 h after self-pollination. JZ\_0.5 and JZ\_2 is that the styles of ‘Jinzhuli’ were collected at 2 h after self-pollination.
(2) Positive values represent up-regulated gene and Negative values represent the down regulated genes. FDR-corrected $P \leq 0.0005$

**Figure 8.**
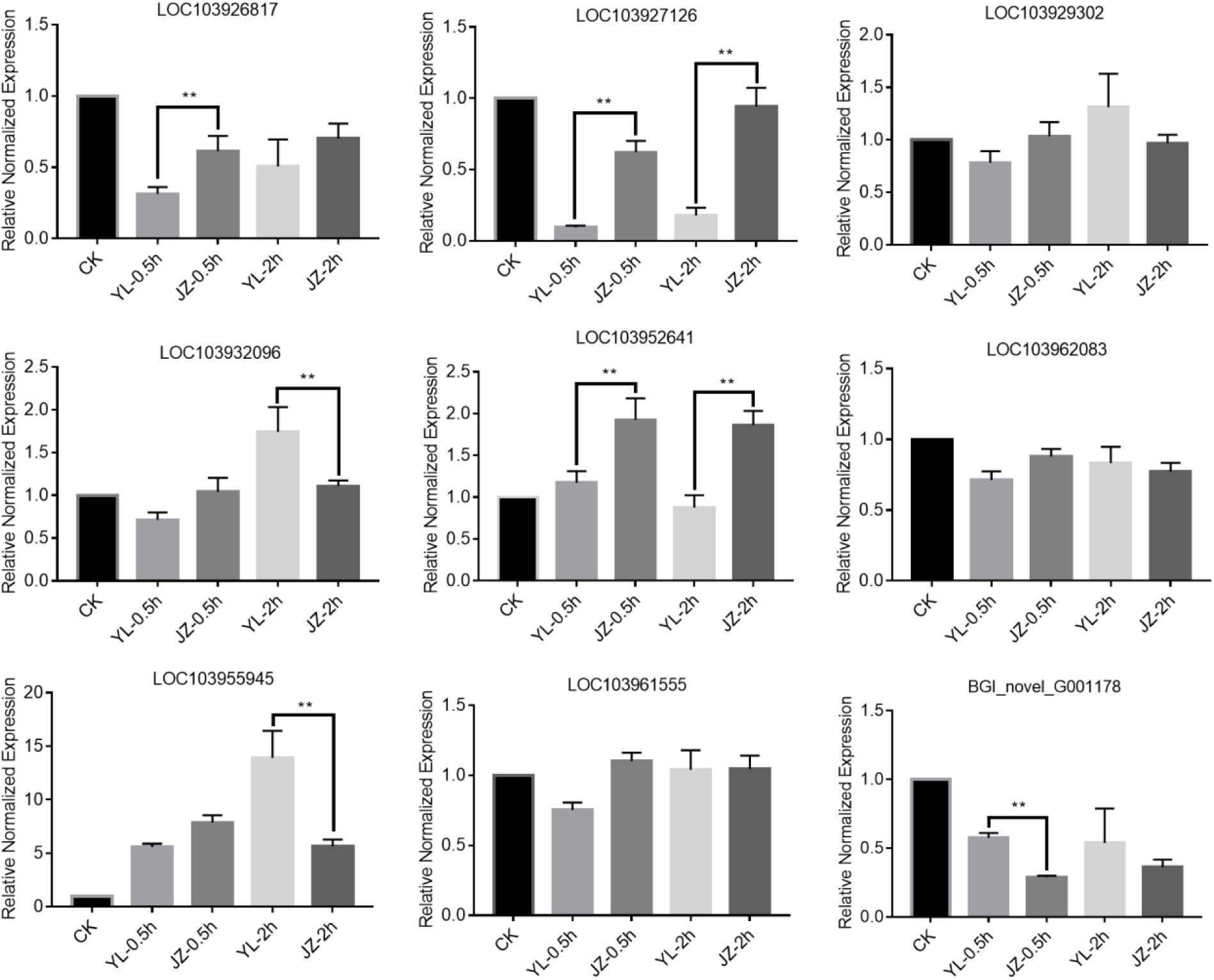
Expression of 10 genes about ATPase in styles measured by quantitative PCR. Comparison of the expression of 10 genes about ATPase during compatible and incompatible pollination. The 10 genes were selected according to the high-throughput sequencing results. ‘Yali’ styles were collected 0.5 and 2 h after self-pollination (YL_0.5h and YL_2h). ‘Jinzhuili’ styles were collected 0.5 and 2 h after self-pollination (JZ_0.5h and JZ_2h). Non-pollinated ‘Yali’ styles were collected after 0.5 h without pollination as the control (YL_CK). The data are presented as the means ± standard error. ** Highly significant data (P < 0.01).

#### Vesicle

Geitmann et al. (2004) suggested that dramatic alterations to the morphology of the mitochondria, Golgi bodies, and endoplasmic reticulum occur within 1 h of SI induction in *Papaver rhoeas* (Geitmann *et al.*, 2004). S-RNase internalization takes place via a membrane/cytoskeleton-based Golgi vesicle system, which can affect self-incompatibility in apple (*Malus domestica*) (Meng *et al.*, 2014). In Brassicaceae, vesicle secretion at the pollen contact site in the stigmatic papilla promotes pollen hydration and pollen tube entry into the stigma through a basal signalling pathway (Safavian & Goring, 2013). The activation of allele-specific (S locus receptor kinase) SCR-SRK (S locus cysteine-rich protein) interactions arrests self-incompatible pollen germination, and the downstream SRK signalling pathway disrupts vesicle secretion from the simultaneously activated basal pollen response pathway (Doucet *et al.*, 2016). These observations indicate that both vesicles in the stigma cells and vesicles in the pollen tubes are involved in the self-incompatibility response. Based on the RNA-Seq results, we selected nine Golgi and vesicle-associated genes for quantitative PCR verification (Table 6). The results of the quantitative expression of these genes were inconsistent with the results of the high-throughput sequencing (Figure 9). Moreover, most of the gene expression was not different when compatibility pollination was compared with incompatible pollination (Figure 9). We speculated that the genes related to Golgi and vesicles are not involved in the mutual recognition of the pollen and stigma at the beginning of pollination.

**Table 6.**
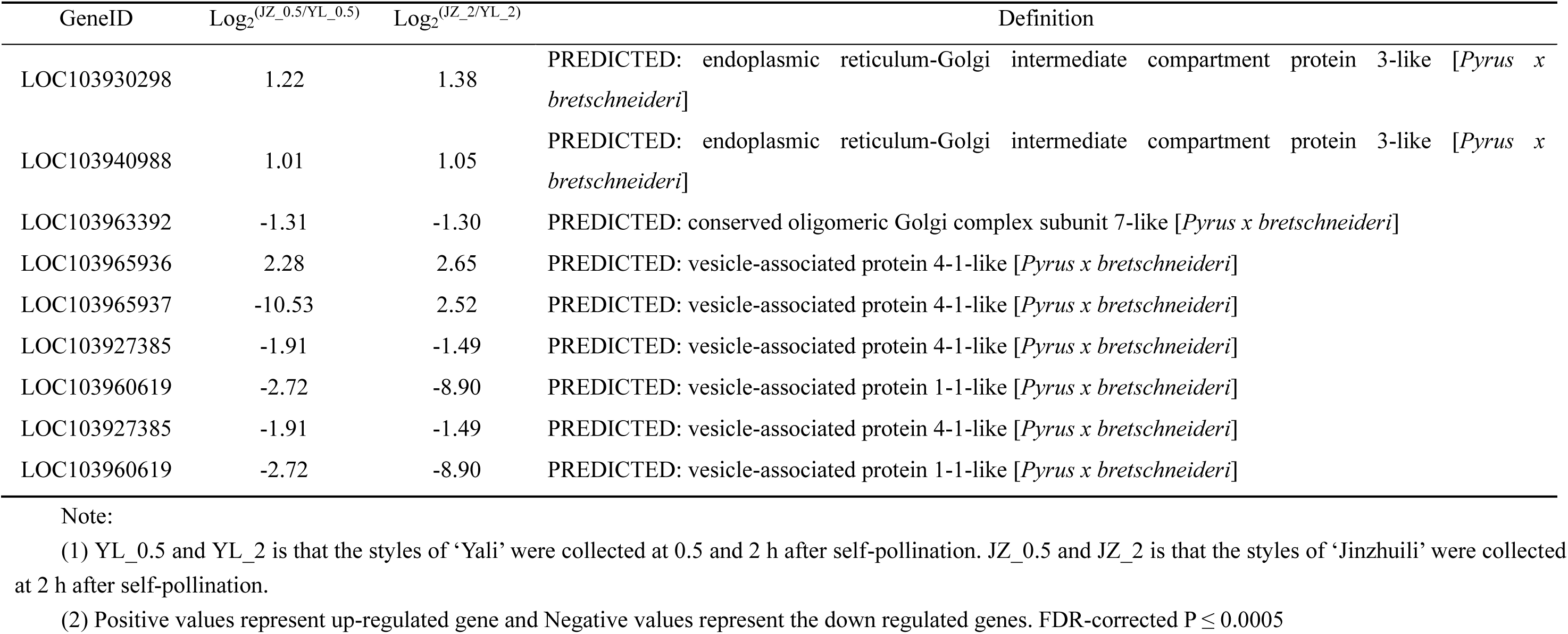
The differences in the expression of 9 genes about Golgi and vesicle and their definitions

**Figure 9.**
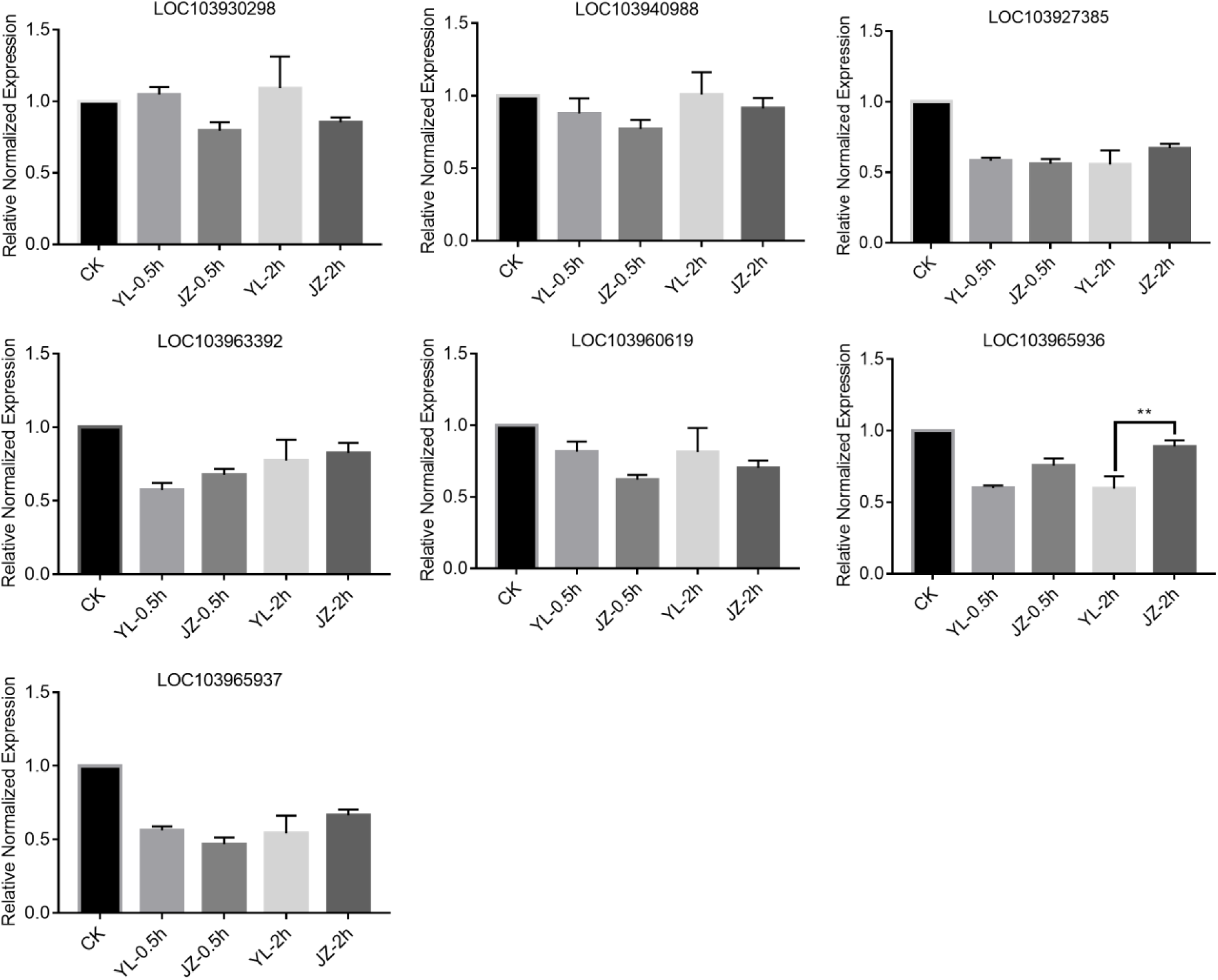
Expression of 9 genes about Golgi and vesicle in styles measured by quantitative PCR. Comparison of the expression of 9 genes about Golgi and vesicle during compatible and incompatible pollination. The 9 genes were selected according to the high-throughput sequencing results. ‘Yali’ styles were collected 0.5 and 2 h after self-pollination (YL_0.5h and YL_2h). ‘Jinzhuili’ styles were collected 0.5 and 2 h after self-pollination (JZ_0.5h and JZ_2h). Non-pollinated ‘Yali’ styles were collected after 0.5 h without pollination as the control (YLCK). The data are presented as the means ± standard error. * * Highly significant data (P < 0.01).

#### SI-induced programmed cell death (PCD)

SI involves highly specific interactions during pollination, resulting in the rejection of incompatible (self) pollen. PCD is an important mechanism for destroying cells in a precisely regulated manner. SI in field poppy (*Papaver rhoeas*) triggers PCD in incompatible pollen (Geitmann *et al.*, 2004; Wilkins *et al.*, 2014). In the self-incompatibility of *Pyrus*, arresting incompatible pollen tube growth requires mechanisms that are partly similar to the SI response of poppy (Wang & Zhang, 2011). In the process of GSI in *Pyrus*, the S-RNase in the style enters the pollen tube and induces PCD in the incompatible pollen tube (Chen *et al.*, 2018; Wang *et al.*, 2010a). There were only two genes related to PCD according to the high-throughout sequencing data, and the difference in expression of the two genes reached significant levels (*P*<0.05) only 2 h after pollination (Table 7). Based on the qt-PCR analysis, no significant differences in the expression of these two genes were observed between the compatible pollination and incompatible pollination at 0.5 or 2 h, respectively (Figure 10). These findings suggest that the pollen tubes were not induced by PCD at the initial stage of incompatibility pollination.

**Table 7.** The differences in the expression of 2 genes about programmed cell death and their definitions.

| GeneID | $\text{Log}_2^{(\text{JZ}_{0.5}/\text{YL}_{0.5})}$ | $\text{Log}_2^{(\text{JZ}_2/\text{YL}_2)}$ | Definition |
| --- | --- | --- | --- |
| LOC103953332 |  | -1.27 | PREDICTED: programmed cell death protein 2 [*Pyrus x bretschneideri*] |
| LOC103926760 |  | 1.19 | PREDICTED: programmed cell death protein 4-like [*Pyrus x bretschneideri*] |

Note:
(1) YL\_0.5 and YL\_2 is that the styles of ‘Yali’ were collected at 0.5 and 2 h after self-pollination. JZ\_0.5 and JZ\_2 is that the styles of ‘Jinzhuli’ were collected at 2 h after self-pollination.
(2) Positive values represent up-regulated gene and Negative values represent the down regulated genes. FDR-corrected $P \leq 0.0005$

**Figure 10.**
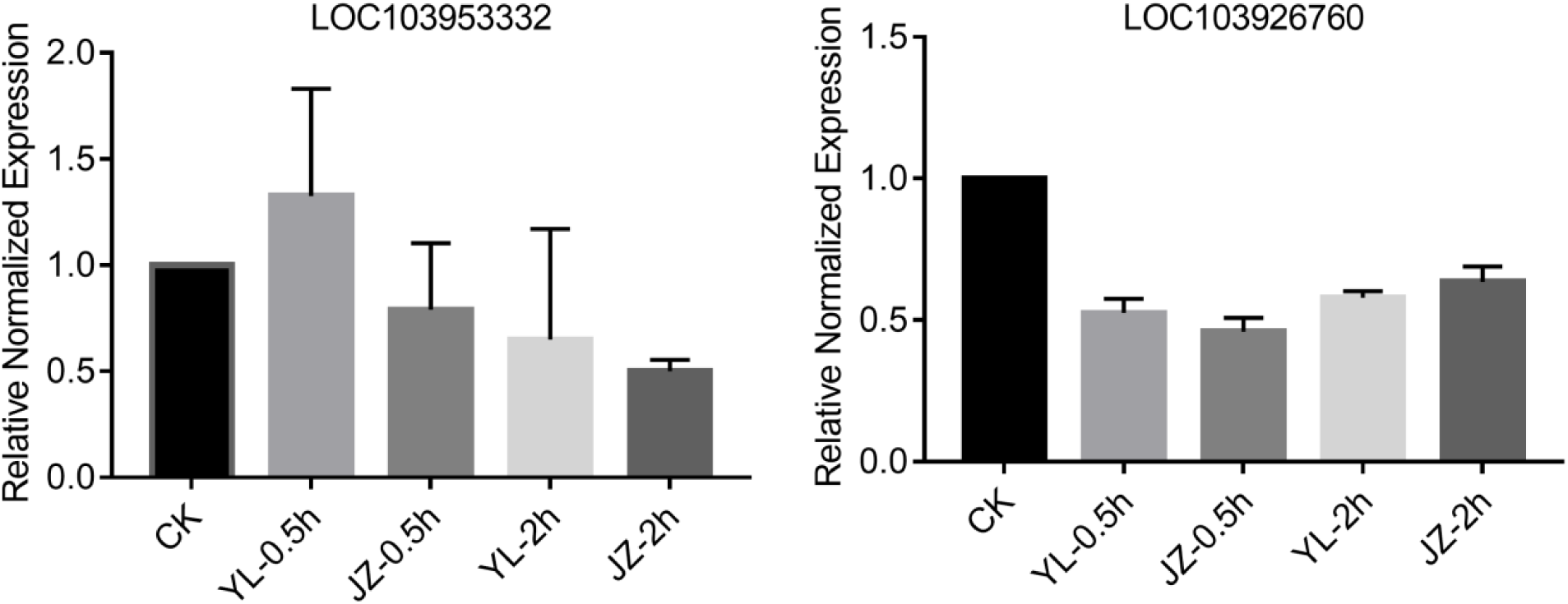
Expression of 2 genes about about programmed cell death in styles measured by quantitative PCR. Comparison of the expression of 2 genes about programmed cell death during compatible and incompatible pollination. The blank cell indicates no difference. The 2 genes were selected according to the high-throughput sequencing results. ‘Yali’ styles were collected 0.5 and 2 h after self-pollination (YL_0.5h and YL_2h). ‘Jinzhuili’ styles were collected 0.5 and 2 h after self-pollination (JZ_0.5h and JZ_2h). Non-pollinated ‘Yali’ styles were collected after 0.5 h without pollination as the control (YLCK). The data are presented as the means ± standard error. * * Highly significant data (P < 0.01).

## Discussion

Apical pollen tubes usually possess a prominent region that includes secretory vesicles, mitochondria, Golgi dictyosomes, and the endoplasmic reticulum (Hepler, 2015). The area contains a large quantity of these subcellular organelles, which is associated with faster and healthier pollen tube growth. Pollen tube growth on the stigma depends on pollen-expressed proteins of the LRX family (Mecchia *et al.*, 2017). In this study, we explored the differences in gene expression between these subcellular organelle-associated genes at the beginning of compatible pollination and incompatible pollination. According to the results of the RNA-Seq and quantitative PCR, once the compatible pollen and incompatible pollen had come into contact with the stigma, genes related to pollen tube growth regulation were significantly upregulated. We previously reported on calcium- and phospholipase C-related genes in the pollen tube that were also significantly differentially expressed in the early stage following pollination (Qu *et al.*, 2017; Qu *et al.*, 2016). All these factors indicate that the gametophytic self-incompatibility response of *Pyrus* has already been initiated after pollination, rather than when the pollen tube enters the style.

The ability to recognize the difference between “self” and “nonself” at the cellular level appears to be dependent on the interactions with the surface components of the organism that come into contact with the plant (Austin & Ballaré, 2014; Lamport *et al.*, 2018; Sequeira, 1978). Plants use recognition mechanisms to reject most of the microorganisms with which they interact (Dresselhaus *et al.*, 2016). Unsurprisingly, the fertilization process in plants uses similar recognition mechanisms to reject foreign pollen and accept pollen from the appropriate species; even the plant’s own pollen may not be able to germinate or penetrate the stigmatic surfaces in those instances where self-incompatibility systems force cross-fertilization (Hodgkin *et al.*, 1988). It is strange why the expression of defensin and resistance protein genes was high at the beginning of compatible pollination, and why the compatible pollen germinates early and grows fast on the stigma. In compatible pollination, the mutual recognition of the pollen and stigma is similar to the plant immune response (Wang *et al.*, 2010b). The substances induced by this immune reaction have a promoting effect on pollen hydration, germination, growth, and orientation (Chae & Lord, 2011). A series of biochemical and genetic studies on plantacyanin and LTP as well as defensin-like proteins have revealed their key role in pollen tube guidance during the fertilization of compatible angiosperms. A series of biochemical and genetic studies on plantacyanin and LTPs and defensin-like proteins have revealed their pivotal roles in pollen tube guidance during compatible angiosperm fertilization (Covey *et al.*, 2010). Although some substances have inhibitory effects on pollen tubes, such as RALF produced by tomato pollen tubes, the activity of RALF depends on LRR. However, RALF inhibits pollen tube growth within 20 minutes. After 20 minutes, the pollen tube grows to a certain length, following which the inhibition disappears (Covey *et al.*, 2010). RALF can also increase the intracellular calcium concentration, which may contribute to the germination and growth of pollen (Covey *et al.*, 2010). In one study, calcium-transporting ATPase gene expression was observed in a compatibility system. In contrast, the interspecific pollination of *A. thaliana* significantly upregulated defensins and inhibited pollen tube growth (Mondragon *et al.*, 2017). An explanation for this inconsistency with our results may be due to the different sampling times after pollination. According to Figure 3, the expression of the defensin gene under incompatible pollination was higher than under compatible pollination at 2 h.

Since the pollen tube can grow to approximately ⅓ of the full length of the style (Wang *et al.*, 2017), which is required at 8 h after self-pollination, it is considered that self-incompatibility in Rosaceae is a stylar event rather than a stigmatic event. Thus far, studies on Rosaceae self-incompatibility have mainly focused on how the S-RNase of the style recognizes self and nonself pollen after pollination. However, at the early stage of pollination, the difference in gene expression between compatibility and incompatibility is highly significant with regards to the pollen-specific expression of the LRR gene, resistance, and defensin genes. Moreover, the interaction between pollen and stigma is similar to the interaction between plants and pathogens only under the condition of compatibility pollination. Therefore, in order to study the self-incompatibility response of *Pyrus*, it is also necessary to study the mutual recognition between cells and between the cell and the signal transduction process as the pollen falls on the stigma.

## Methods

### Plant materials

The self-compatible pear variety ‘Jinzhuili’ was a naturally occurring bud mutant from ‘Yali’, a leading Chinese native cultivar with typical GSI. ‘Jinzhuili’ had lost the pollen SI function and that its style behaved normally in SI response, similar to that of the wild type ‘Yali’ (Wu *et al.*, 2013a). We collected the styles of more than 100 flowers of the ‘Yali’ and ‘Jinzhui’ at 0.5, 2 h after self-pollination respectively and the styles of ‘Yali’ without pollination as control. Respectively marked as YL_0.5h, YL_2h, JZ_0.5h, JZ_2h and YL_CK. All of the harvested samples were immediately deep-frozen in liquid nitrogen and stored in liquid nitrogen before use.

### RNA isolation, validation, and RNA-seq library preparation

RNA was isolated from each of the above style. Therefore, a total of three RNA bulks representing of styles of the self-incompatibility and non-pollination and the self-compatibility were available for transcriptome sequencing analysis. Total RNA was extracted with TRIzol reagent (Life Technologies, Carlsbad, CA, USA) according to the manufacturer’s instruction and treated with RNase-free DNase I (Takara Biotechnology, Dalian, China). An Agilent 2100 Bioanalyzer (Agilent, Santa Clara, CA, USA) was then utilized to confirm RNA integrity based on a minimum RNA integrated threshold value of eight. Poly(A) mRNA was isolated with oligo-dT beads and then treated with fragmentation buffer. The cleaved RNA fragments were then transcribed into first-strand cDNA using reverse transcriptase and random hexamer primers. This was followed by second strand cDNA synthesis using DNA polymerase I and RNaseH. The double-stranded cDNA was further subjected to end-repair using T4 DNA polymerase, Klenow fragment, and T4 Polynucleotide kinase followed by a single nucleotide “A” base addition using Klenow exo2 polymerase. The cDNAs were then ligated with adapters using T4 DNA ligase. Adaptor ligated fragments were subjected to agarose gel electrophoresis and cDNA fragments within the desired size range (304 ~ 374 bps) were excised from the gel. PCR was performed to amplify these fragments. The quality of the cDNA fragments was validated using an Agilent 2100 Bioanalyzer and a StepOnePlus Real-Time PCR System (Life Technologies, Carlsbad, CA, USA), after which the cDNA library was sequenced using a flow cell HiSeq2000 sequencer (Illumina, San Diego, CA, USA).

### Pathway analysis

Pathway analysis was used to determine the significant pathway(s) of the DEGs according to the KEGG database. Fisher’s exact test was used to select significant pathways, and the threshold of significance was defined according to the *P* value and FDR (Draghici *et al.*, 2007; Kanehisa *et al.*, 2004; Yi *et al.*, 2006).

### Quantitative real-time RT-PCR to determine gene expression in style

To estimate mRNA expression levels for genes, qRT-PCR analysis was performed. Style was got and conserved in liquid nitrogen after manual pollination for 0.5, 1 and 2 h. Total RNA was extracted with TRIzol regent (Invitrogen), and the first-strand cDNA was synthesized from 2 μg of total RNA using an Omniscript RT kit (Qiagen). All primers were synthesized at HUADA (The Central Facility of the Institute for Molecular Biology and Biotechnology, McMaster University). Gene expression data were normalized to Actin levels and expressed as ratios to the relevant control (assigned a value of 1).

### Graphics software

DE genes with FDR < 0.005 for comparisons between groups of samples. Statistical mapping using GraphPad Prism 7.0 software.

## Acknowledgments

We would like to thank all scientists for helping us to improve the manuscript. The work was supported by Doctoral Fund of Qingdao Agricultural University, experts working in the fruit system of Shandong Province.

